# CCL28 modulates neutrophil responses during infection with mucosal pathogens

**DOI:** 10.1101/2021.03.19.436197

**Authors:** Araceli Perez-Lopez, Steven Silva, Michael H. Lee, Nicholas Dillon, Stephanie L. Brandt, Romana R. Gerner, Karine Melchior, Gregory T. Walker, Victor A. Sosa-Hernandez, Rodrigo Cervantes-Diaz, Sandra Romero-Ramirez, Jose L. Maravillas-Montero, Sean-Paul Nuccio, Victor Nizet, Manuela Raffatellu

## Abstract

The chemokine CCL28 is highly expressed in mucosal tissues, but its role during infection is not well understood. Here we show that CCL28 promotes neutrophil accumulation to the gut of mice infected with *Salmonella* and to the lung of mice infected with *Acinetobacter*. Neutrophils isolated from the infected mucosa expressed the CCL28 receptors CCR3 and, to a lesser extent, CCR10, on their surface. The functional consequences of CCL28 deficiency were different between the two infections, as *Ccl28*^-/-^ mice were highly susceptible to *Salmonella* gut infection, but highly resistant to otherwise lethal *Acinetobacter* lung infection. *In vitro*, unstimulated neutrophils harbored pre-formed intracellular CCR3 that was rapidly mobilized to the cell surface following phagocytosis or inflammatory stimuli. Moreover, CCL28 stimulation enhanced neutrophil antimicrobial activity, production of reactive oxygen species, and formation of extracellular traps, all processes that were largely dependent on CCR3. Consistent with the different outcomes in the two infection models, neutrophil stimulation with CCL28 boosted the killing of *Salmonella* but not of *Acinetobacter*. CCL28 thus plays a critical role in the immune response to mucosal pathogens by increasing neutrophil accumulation and activation, which can enhance pathogen clearance but also exacerbate disease depending on the mucosal site and the infectious agent.

## Introduction

Chemokines comprise a family of small chemoattractant proteins that play important roles in diverse host processes including chemotaxis, immune cell development, and leukocyte activation (1–3). The chemokine superfamily includes 48 human ligands and 19 receptors, commonly classified into subfamilies (CC, CXC, C, and CX_3_C) depending on the location of the cysteines in their sequence (4, 5). Four chemokines predominate in mucosal tissues: CCL25, CCL28, CXCL14, and CXCL17 (6).

CCL28, also known as Mucosae-associated Epithelial Chemokine (MEC), belongs to the CC (or β-chemokine) subclass, and is constitutively produced in mucosal tissues including the digestive system, respiratory tract, and female reproductive system (7). Although best studied for its homeostatic functions, CCL28 can also be induced under inflammatory conditions and is thus considered a dual function (homeostatic and inflammatory) chemokine (7).

CCL28 signals via two receptors: CCR3 and CCR10 (8). During homeostasis in mice, CCL28 provides a chemotactic gradient for CCR10⁺ B and T cells and guides the migration of CCR10⁺ IgA plasmablasts to the mammary gland and other tissues (7, 9, 10). In a disease context, CCL28 has been best studied in allergic airway inflammation. High CCL28 levels are present in airway biopsies from asthma patients (11), and CCR3^+^ and CCR10^+^ cells are recruited to the airways in a CCL28-dependent fashion in murine asthma models (12, 13).

In the human gut, CCL28 is upregulated during inflammation of the gastric mucosa in *Helicobacter pylori*-infected patients (14) and in the colon of patients with ulcerative colitis, a prominent form of inflammatory bowel disease (15, 16). In the mouse gut, CCL28 production is increased in the dextran sulfate sodium (DSS) model of colitis (10). Epithelial cells are an important source of CCL28 (15, 16), and its expression can be induced by stimulation of cultured airway or intestinal epithelial cells with the proinflammatory cytokines IL-1ɑ, IL-1β, or TNFɑ, or following *Salmonella* infection of cultured HCA-7 colon carcinoma cells (16).

Collectively, a variety of studies have postulated that CCL28 is an important chemokine in inflammatory diseases, ranging from asthma to ulcerative colitis and during the immune response to infection. Yet, CCL28 function remains understudied, largely because *Ccl28*^-/-^ mice have only recently been described (9, 10). Here, we investigate the function and underlying mechanism of CCL28 during the mucosal response to infection.

By comparing infection in *Ccl28*^-/-^ mice and their wild-type littermates, we discovered a key role for CCL28 in promoting neutrophil accumulation to the gut during infection with *Salmonella enterica* serovar Typhimurium (STm) and to the lung during infection with multidrug-resistant *Acinetobacter baumannii* (Ab). Neutrophils isolated from the infected mucosal sites harbored CCL28 receptors, particularly CCR3, on their surface. *In vitro*, CCR3 was stored intracellularly and was mobilized to the surface upon neutrophil stimulation with proinflammatory molecules or in response to phagocytosis. Neutrophil stimulation of CCL28 resulted in enhanced neutrophil antimicrobial activity against STm, increased production of reactive oxygen species (ROS), and enhanced formation of neutrophil extracellular traps (NETs), all processes that help control infection but also cause extensive tissue damage. We conclude that CCL28 plays a previously unappreciated role in the innate immune response to mucosal pathogens by regulating neutrophil accumulation and activation.

## Results

### CCL28-mediated responses limit *Salmonella* gut colonization and systemic dissemination

We investigated CCL28 activity during gastrointestinal infection with the enteric pathogen *Salmonella enterica* serovar Typhimurium (STm) by using the well-established streptomycin-treated C57BL/6 mouse model of colitis (17). At 96h post-infection with STm, we observed a ∼4-fold increase of CCL28 by ELISA analysis of feces from wild-type mice relative to uninfected controls (**Fig. 1A**). In a preliminary study, we found that *Ccl28^-/-^* mice infected with STm exhibited increased lethality compared to their wild-type littermates beginning at 24h post-infection (9). As such, we enumerated STm colony-forming units (CFU) in tissues at 72h post-infection, instead of the frequently studied 96h endpoint. We recovered significantly higher STm CFU from the gastrointestinal tract (cecum content, Peyer’s patches), the mesenteric lymph nodes, and systemic sites (bone marrow and spleen) of *Ccl28^-/-^* mice vs. wild-type littermates (**Fig. 1B**), demonstrating that the chemokine is an essential component of host defense against STm in this colitis model. When bypassing the gut by infecting mice with STm via the intraperitoneal route, we also observed a ∼4-fold increase of CCL28 in the serum (**Suppl. Fig. 1A**); however, we recovered equal numbers of STm CFU in the spleen, liver, and blood of wild-type and *Ccl28^-/-^* mice (**Suppl. Fig. 1B**). Thus, CCL28 helps the host to control STm infection at its point of origin in the gut mucosa, prompting us to further investigate the underlying mechanisms of CCL28-mediated mucosal protection.

**Figure 1.**
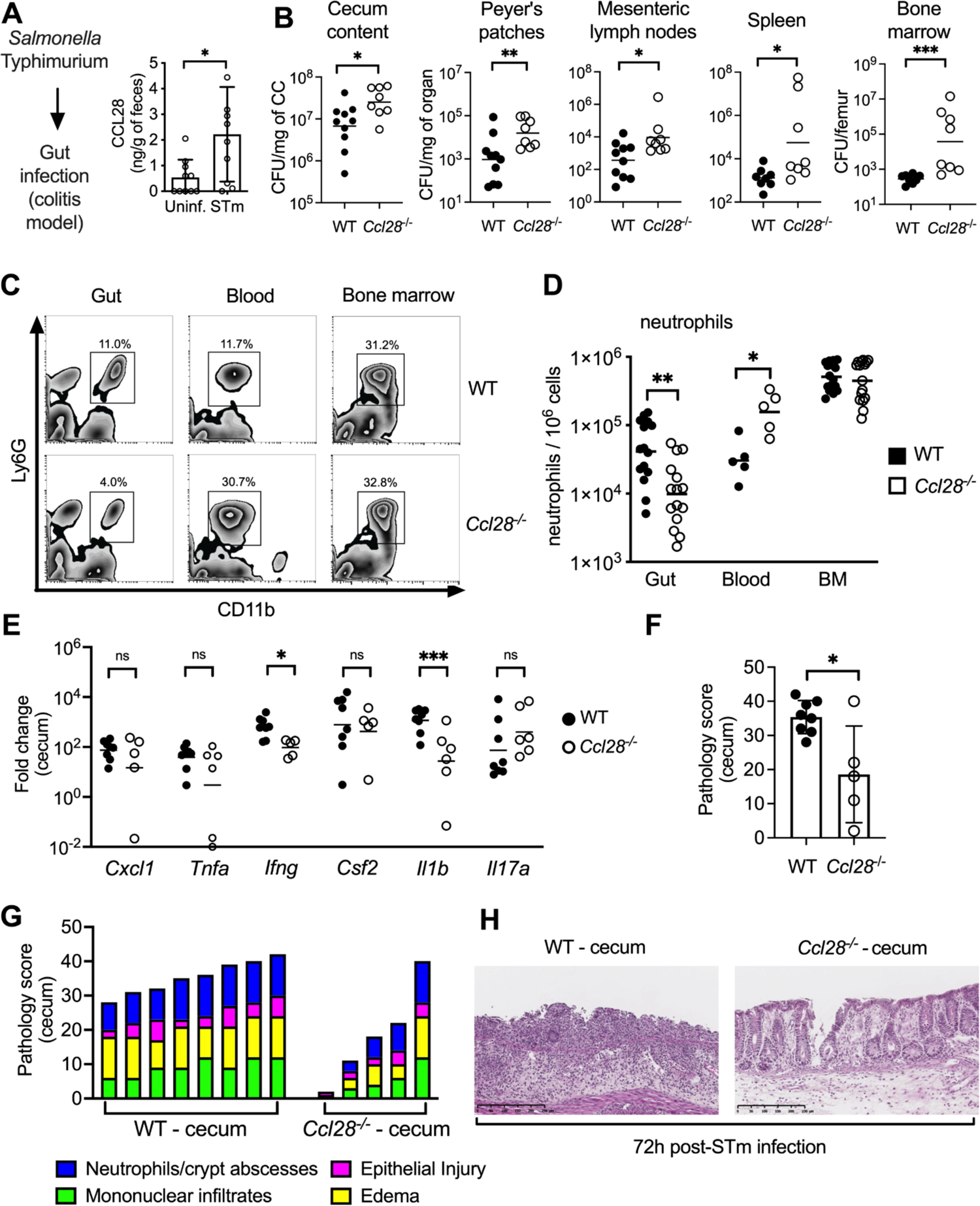
CCL28 confers protection during *Salmonella* colitis and promotes neutrophil accumulation in the gut. (**A**) For the colitis model, wild-type mice were gavaged with streptomycin 24h prior to oral infection with *S. enterica* serovar Typhimurium (STm). At 96h post-infection, CCL28 in feces was quantified by ELISA. Data shown comprise two independent experiments (uninfected, n=10; STm, n=10). Bars represent the mean ± SD. (**B**) CFU in cecum content, Peyer’s patches, mesenteric lymph nodes, spleen, and bone marrow were determined at 72h post-infection of wild-type (WT, black circles) and *Ccl28^-/-^* (white circles) littermate mice. Data shown comprise three independent experiments (WT, n=10; *Ccl28^-/-^*, n=8). Some of the spleen data points were published as a preliminary characterization in Burkhardt et al. (Ref 9) and are combined with the new dataset. Bars represent the geometric mean. (**C**) Representative contour plots of neutrophils (CD11b^+^ Ly6G^+^ cells; gated on live, CD45^+^ cells) obtained from the gut, blood, and bone marrow of STm-infected WT or *Ccl28^-/-^* mice, as determined by flow cytometry at 48h post-infection. (**D**) Frequency of neutrophils per million live CD45+ cells obtained from the gut, blood, and bone marrow of STm-infected WT (black bars) or *Ccl28^-/-^* mice (white bars). Data shown comprise three independent experiments (WT, n=15; *Ccl28^-/-^*, n=15; blood was collected from n = 5 per group). Bars represent the mean ± SD. (**E**) Relative expression levels (qPCR) of *Cxcl1* (CXCL1), *Tnfa* (TNFα), *Ifng* (IFNγ), *Csf2* (GM-CSF), *Il1b* (IL-1β), and *Il17a* (IL-17A) in the cecum of WT (black circles, n=8) or *Ccl28^-/-^* mice (white circles, n=6). Bars represent the geometric mean. Data shown comprise three independent experiments. (**F-H**) Histopathological analysis of cecum collected from STm-infected WT or *Ccl28^-/-^* mice (WT, n=8; *Ccl28^-/-^*, n=5). (**F**) Sum of the total histopathology score, (**G**) histopathology scores showing the analyzed parameters, and (**H**) H&E-stained sections from one representative animal for each group (200X). (**F**) Bars represent the mean ± SD. Mann-Whitney U was used for all datasets where statistical analysis was performed. A significant change relative to WT controls is indicated by \**p* ≤ 0.05, \*\**p* ≤ 0.01, \*\*\**p* ≤ 0.001. ns = not significant.

### CCL28 promotes neutrophil accumulation to the gut during *Salmonella* infection

CCL28 has direct antimicrobial activity against some bacteria (e.g., *Streptococcus mutans* and *Pseudomonas aeruginosa*) and fungi (e.g., *Candida albicans*) (18). However, concentrations of CCL28 up to 1μM did not substantially inhibit wild-type STm, whereas the chemokine produced multilog-fold CFU reductions in *Escherichia coli* K12 or an STm Δ*phoQ* mutant, which is more susceptible to antimicrobial peptides than STm wild-type (19), at similar concentrations (**Suppl. Fig. 1C**). Thus direct antimicrobial activity of CCL28 is unlikely to explain the lower STm colonization in wild-type mice in comparison to *Ccl28^-/-^* mice.

During homeostasis, CCL28 exhibits chemotactic activity in the gut mucosa towards CD4^+^ and CD8^+^ T cells and IgA-producing B cells (7, 9, 10). However, B and T cell numbers in the intestine of wild-type and *Ccl28^-/-^* mice were similar during homeostasis (**Suppl. Fig. 2A** and **2C**) and 48h after STm infection (**Suppl. Fig. 2B** and **2D**), indicating that the chemokine’s protective role is likely independent of its B or T cell chemotactic activity. Neutrophils are a crucial component of the host response to STm (reviewed in (20)), and neutropenia increases the severity of infection with non-typhoidal *Salmonella* in both mice and humans (21–24). Strikingly, we isolated ∼50% fewer neutrophils (CD11b⁺ Ly6G⁺ cells) from the gut of *Ccl28^-/-^* mice 48h after STm infection relative to their wild-type littermates (**Fig. 1C, D**). Commensurate with this finding, neutrophil counts in the blood of infected *Ccl28^-/-^* mice were increased compared to wild-type mice (**Fig. 1C, D**), indicating a defect in recruitment of circulating neutrophils to the site of infection. Neutrophil counts in the bone marrow were similar between wild-type and *Ccl28^-/-^* animals (**Fig. 1C, D**), excluding a defect in granulopoiesis. Likewise, bone marrow and blood neutrophil counts were similar in wild-type and *Ccl28^-/-^* mice under homeostatic conditions (**Suppl. Fig. 2E, F**) and neutrophils were not found in uninfected gut tissue (data not shown). Thus, CCL28 promotes neutrophil accumulation to the gut during STm infection by a mechanism that transpires after bone marrow neutrophil production and their egress into the blood circulation.

**Figure 2.**
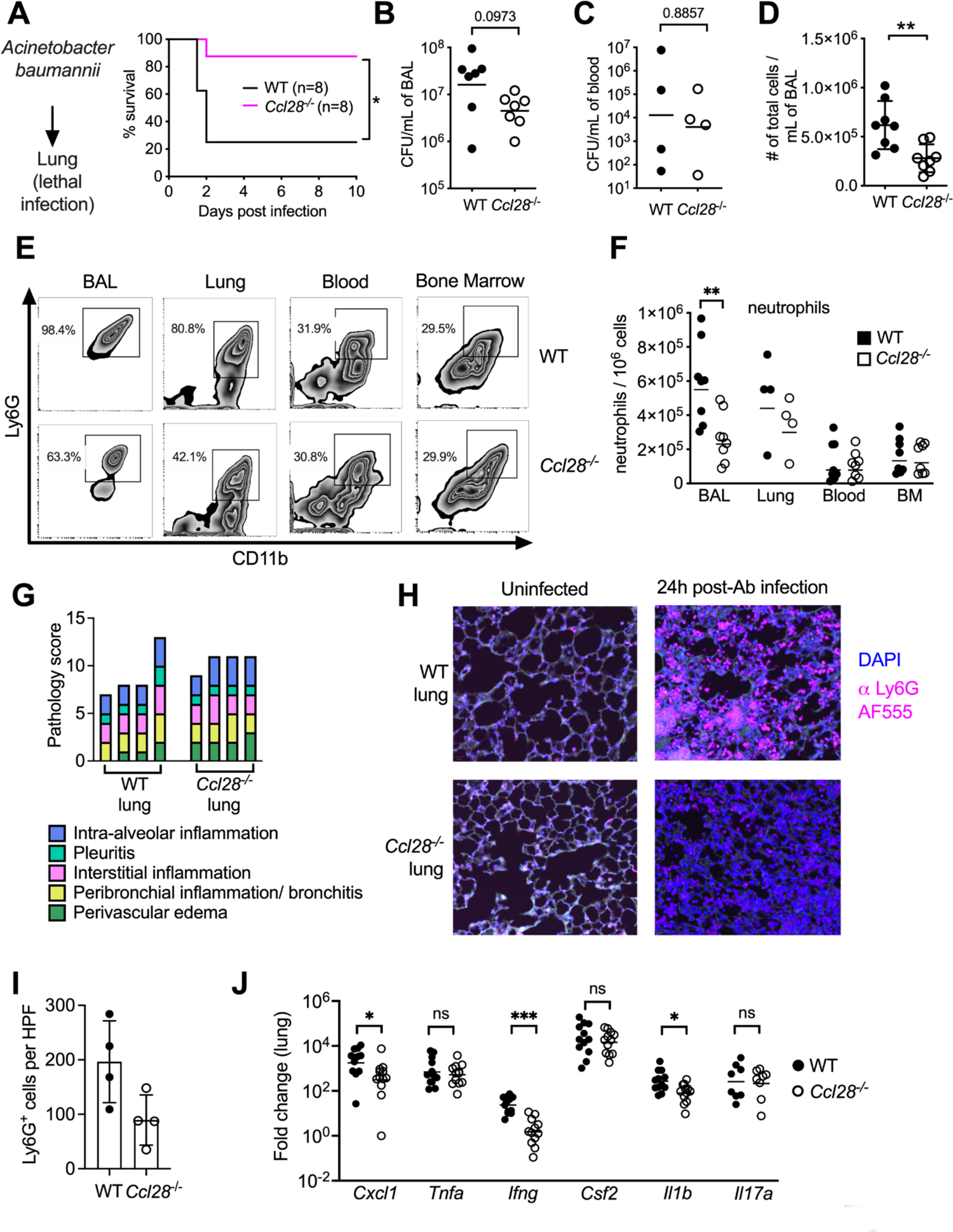
Absence of CCL28 confers protection in a lethal *Acinetobacter* pneumonia model. (**A**) WT mice (black line) and *Ccl28^-/-^* mice (magenta line) were intratracheally infected with *Acinetobacter baumannii* (Ab) and their survival was determined for 10 days. Data shown comprise two independent experiments (WT, n=8; *Ccl28^-/-^*, n=8). (**B-F**) WT and *Ccl28^-/-^* mice were compared 24h post-infection with Ab. (**B, C**) Ab CFU in (**B**) BAL (bronchoalveolar lavage) fluid or (**C**) blood in WT mice (black circles) and *Ccl28^-/-^* mice (white circles). (**D**) The number of total host cells per mL of BAL, as determined by flow cytometry. (WT, n=8; *Ccl28^-/-^*, n=8) (**E, F**) Representative (**E**) contour plots of neutrophils (CD11b^+^ Ly6G^+^ cells; gated on live, CD45^+^ cells) and (**F**) frequency of neutrophils obtained from the BAL, lung, blood, and bone marrow of Ab-infected WT or *Ccl28^-/-^* mice, as determined by flow cytometry. Data comprise two independent experiments (WT, n=8; *Ccl28^-/-^*, n=8; lung was collected from n = 4 per group). Bars represent (**B, C)** the geometric mean or (**D, F**) the mean ± SD. (**G**) Histopathological analysis of lungs from WT and *Ccl28^-/-^* mice infected with Ab at 24 h post-infection. Each bar represents an individual mouse. (**H**) Representative immunofluorescence image of lungs from WT and *Ccl28^-/-^* mice, uninfected or infected with Ab stained for the neutrophil marker Ly6G (magenta). DAPI (blue) was used to label nuclei. (**I**) Quantification of Ly6G^+^ cells per high-power field (HPF) from immunofluorescence images of lungs from WT mice (n=4) and *Ccl28^-/-^* mice (n=4). Bars represent the mean ± SD. (**J**) Relative expression levels (qPCR) of *Cxcl1* (CXCL1), *Tnfa* (TNFα), *Ifng* (IFNγ), *Csf2* (GM-CSF), *Il1b* (IL-1β), and *Il17a* (IL-17A) in the lung of WT (black circles, n=12) or *Ccl28^-/-^* mice (white circles, n=12) infected with Ab. Bars represent the geometric mean. Data shown comprise three independent experiments. Mann-Whitney U was used for all datasets where statistical analysis was performed. A significant change relative to WT controls is indicated by \**p* ≤ 0.05, \*\**p* ≤ 0.01, \*\*\**p* ≤ 0.001. ns = not significant.

### Gut proinflammatory gene expression and tissue pathology are reduced in *Ccl28*^-/-^ mice infected with STm

Neutrophils can mediate inflammation by directly producing proinflammatory molecules or by engaging in crosstalk with other cells (25). We evaluated the expression of genes encoding proinflammatory cytokines in the cecum of *Ccl28^-/-^* mice and their wild-type littermates 72h after STm infection. IFNγ and IL-1β gene transcripts were significantly higher in the cecum of infected wild-type mice vs. *Ccl28^-/-^* mice, while expression of genes encoding other factors involved in neutrophil recruitment (CXCL1, GM-CSF, IL-17A) or the proinflammatory cytokine TNF-ɑ did not differ significantly (**Fig. 1E**). No difference in the expression of these genes was found between uninfected wild-type mice and *Ccl28^-/-^* mice (data not shown). Consistent with the role of neutrophils as important mediators of inflammation in STm colitis, histopathology at 72h post-infection revealed marked cecal inflammation (including significant neutrophil recruitment) in wild-type mice that was greatly reduced in *Ccl28^-/-^* mice (**Fig. 1F-H**). Thus, by modulating neutrophil accumulation to the infected gut, CCL28 drives inflammatory tissue pathology and colitis during STm infection.

### *Ccl28^-/-^* mice are protected from lethal infection in an *Acinetobacter* pneumonia model

CCL28 is expressed in several mucosal tissues beyond the gut, including the lung (7). To investigate whether CCL28 also promoted neutrophil accumulation and host protection in the lung during infection, we employed a murine *Acinetobacter baumannii (*Ab) pneumonia model (26, 27). Ab is an emerging, frequently multidrug-resistant Gram-negative pathogen that causes potentially lethal nosocomial pneumonia in intensive care unit patients (28). Following intratracheal Ab infection, we observed a striking and unexpected phenotype in the mortality curves of wild-type vs. *Ccl28^-/-^* mice: while 6 out of 8 wild-type littermates (75%) died within 48h, 7 out of 8 of the *Ccl28^-/-^* knockout mice (88%) survived through Day 10 post-infection (**Fig. 2A**). The enhanced resistance of *Ccl28^-/-^* mice relative to their wild-type littermates was not associated with a significant reduction in Ab CFU recovered from bronchoalveolar lavage (BAL) fluid or blood (**Fig. 2B, C**). Thus, opposite to what we observed during STm gut infection, CCL28 did not confer protection during Ab lung infection, but rather exacerbated lethality. *In vitro*, high concentrations (1μM) of CCL28 exhibited direct antimicrobial activity against 5×10^5^ CFU of Ab, but not when higher CFU (5×10^8^/ml) were used as inoculum in the assay (**Suppl. Fig. 1C**). Given that high Ab CFU were recovered in the lung of wild-type mice (**Fig. 2B**), CCL28 does not appear to limit growth of this pathogen *in vivo* even though it exhibits antimicrobial activity *in vitro.* We thus investigated whether there were differences in neutrophil accumulation in the lung between wild-type and *Ccl28^-/-^* mice that could explain the higher lethality of *Ccl28^-/-^* mice during Ab lung infection.

### CCL28 promotes neutrophil accumulation to the lung during *Acinetobacter* infection

Prior studies demonstrated that neutrophils are recruited to the lungs of mice infected with Ab beginning at 4h post-infection, and peak at 24h post-infection (29, 30). As CCL28 contributed to neutrophil recruitment during STm gut infection, we analyzed neutrophil recruitment to the lung mucosa 24h after Ab infection in wild-type and *Ccl28^-/-^* mice. BAL fluid showed a significantly greater cellular infiltrate (largely comprised of neutrophils; CD11b⁺ Ly6G⁺ cells) in wild-type mice than in *Ccl28^-/-^* littermates (**Fig. 2D-F**). A similar higher percentage of neutrophils was identified in lung tissue of wild-type mice compared to *Ccl28^-/-^* mice post-infection, whereas we found no differences in neutrophil numbers in the blood or bone marrow (**Fig. 2E-F**). Although histopathology of lung tissue found substantial lung inflammation in both wild-type and *Ccl28^-/-^* mice after Ab infection (**Fig. 2G**), immunofluorescence analysis revealed remarkable differences in the composition of the cellular infiltrate — lungs from infected wild-type mice showed an abundant infiltrate of neutrophils (Ly6G^+^ cells) that was significantly less in *Ccl28^-/-^* mice (**Fig. 2H-I**).

Comparable to what we observed during STm gut infection, induction of the genes encoding IFNγ and IL-1β was significantly lower in Ab-infected lungs of *Ccl28^-/-^* mice relative to wild-type mice (**Fig. 2J**). By contrast, expression of the *Cxcl1* gene was reduced, whereas the other proinflammatory genes tested (*Il17a*, *Csf2*, *Tnfa*) did not differ between the groups (**Fig. 2J**). CCL28 thus contributes to lung inflammation and neutrophil accumulation during Ab pneumonia, which is consistent with our findings during STm gut infection. We thus sought to further investigate how CCL28 regulates neutrophil responses.

### Gut and BAL neutrophils express receptors CCR3 and CCR10 during infection

CCL28 attracts leukocytes that express at least one of its receptors (CCR3, CCR10). CCR10 is found on T cells, B cells, and IgA-secreting plasma cells, whereas eosinophils express CCR3 (7). Although early studies concluded that CCR3 was absent in neutrophils (31), the receptor was later detected on the surface of neutrophils isolated from patients with chronic inflammation (32). Based on these prior studies and our findings of CCL28-dependent neutrophil accumulation to the gut during gut and lung infection (**Fig. 1, 2**), we performed flow cytometry on single-cell suspensions from tissue isolated from infected mice to evaluate surface expression of CCR3 and CCR10. For mice infected with STm, we analyzed the gut, blood, and bone marrow (**Fig. 3A, B**). While both chemokine receptors were identified on a subset of bone marrow neutrophils (∼36% CCR3, ∼12% CCR10) and blood neutrophils (∼25% CCR3, ∼16% CCR10) during infection, neutrophils expressing these receptors were enriched in the inflamed gut, with ∼80% expressing CCR3 and ∼28% expressing CCR10 (**Fig. 3A**, **B**). In a separate experiment where neutrophils were simultaneously stained for both CCR3 and CCR10, ∼35% of gut neutrophils from infected wild-type mice expressed both receptors (**Suppl. Fig. 3A)**, and infected *Ccl28^-/-^* mice expressed similar levels of these receptors as wild-type mice (**Suppl. Fig. 3B)**.

**Figure 3.**
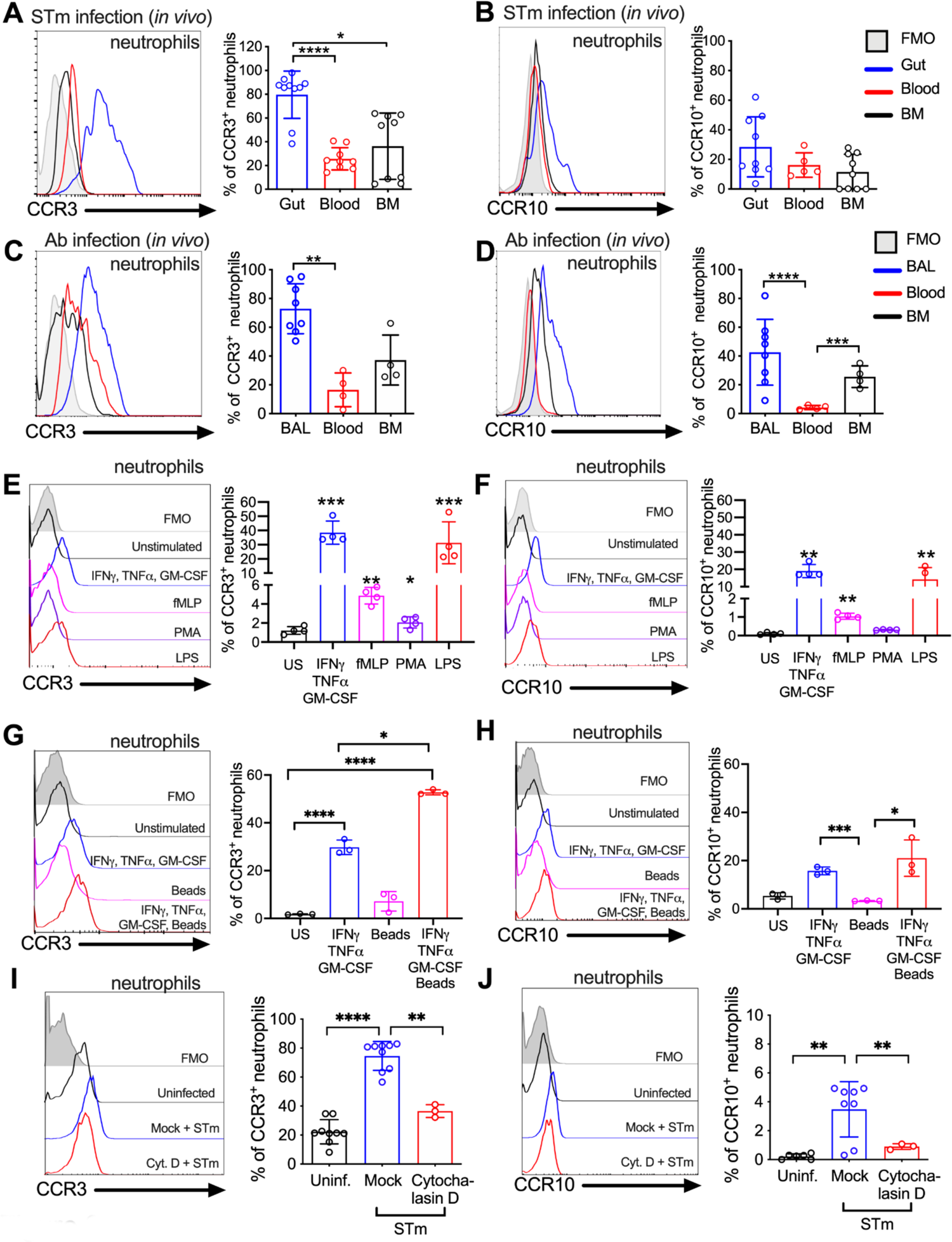
Surface expression of the CCL28 receptors CCR3 and CCR10 on neutrophils from infected tissue, and upon stimulation with proinflammatory stimuli and phagocytosis. (**A-D)** Surface expression of (**A, C**) CCR3 or (**B, D**) CCR10 on neutrophils obtained from (**A, B**) the gut, blood, and bone marrow (BM) 72h post-infection with STm, or (**C, D**) the BAL, blood, and bone marrow 24h post-infection with Ab, analyzed by flow cytometry. Left panels show representative histograms of (**A, C**) CCR3 or (**B, D**) CCR10 expression on the surface of neutrophils (gated on live, CD45^+^ CD11b^+^ Ly6G^+^ cells) from (**A, B**) the gut (blue), blood (red), and bone marrow (BM; black) or (**C, D**) BAL (blue), blood (red), and bone marrow (BM; black). Right panels show the percentage of (**A, C**) CCR3^+^ or (**B, D**) CCR10^+^ neutrophils obtained from (**A, B**) gut, blood, and BM or (**C, D**) BAL, blood, and BM. Data are from two independent experiments. (**E-H**) Uninfected bone marrow neutrophils were unstimulated or treated with the indicated stimuli for 4h. Surface expression of (**E**, **G**) CCR3 and (**F**, **H**) CCR10 on neutrophils was determined by flow cytometry. Left panels show representative histograms of (**E**, **G**) CCR3 or (**F**, **H**) CCR10 surface expression after stimulation with: (**E, F**) cytokines IFNγ + TNFɑ + GM-CSF (blue); fMLP (magenta); PMA, (purple); LPS (red); (**G, H**) cytokines IFNγ + TNFɑ + GM-CSF (blue); beads alone (magenta); cytokines plus beads (red). Right panels show the percentage of (**E, G**) CCR3^+^ or (**F**, **H**) CCR10^+^ neutrophils following stimulation with the indicated stimuli. US = unstimulated. Data shown are representative of two independent experiments. (**I, J**) Bone marrow neutrophils were infected with opsonized STm at a multiplicity of infection (MOI)=10 for 1h with (red) or without (blue) pretreatment with cytochalasin D for 30 min before infection. Surface expression of (**I**) CCR3 or (**J**) CCR10 was determined by flow cytometry. Data are from three independent experiments. Left panels show representative histograms of surface receptor staining on neutrophils, and right panels show the percentages. (**A-J**, right panels) Bars represent the mean ± SD. (**A-D, I, J**) Log-transformed data were analyzed by one-way ANOVA for independent samples, assuming non-equal SD given the differences in the variance among the groups and followed by Dunnett’s multiple comparisons test. (**E-H**) Data were analyzed by one-way ANOVA for paired samples, applying the Greenhouse-Geisser correction given the differences in variance among the groups; (**E, F**) Dunnett’s multiple comparison test was performed to compare the different conditions of stimulation with the control group; (**G, H**) Tukey’s multiple comparison test was performed to compare each condition to each other. Significant changes are indicated by \**p* ≤0.05, \*\**p* ≤0.01, \*\*\**p* ≤0.001, \*\*\*\**p* ≤ 0.0001.

Neutrophils isolated from the BAL of Ab-infected wild-type mice were also positive for CCR3 and CCR10 surface expression, with ∼73% of neutrophils expressing CCR3 (**Fig. 3C**) and ∼43% expressing CCR10 (**Fig. 3D**). In a separate experiment where neutrophils were simultaneously stained for both CCR3 and CCR10, ∼20% of BAL neutrophils from infected wild-type mice expressed both receptors (**Suppl. Fig. 3C)**, and infected *Ccl28^-/-^* mice expressed similar levels of these receptors as wild-type mice (**Suppl. Fig. 3D)**. As predicted, compared to BAL neutrophils, a lower percentage of neutrophils isolated from the blood and the bone marrow of mice infected with Ab were positive for the receptors (**Fig. 3C, D**). Thus, neutrophils from the infected mucosal tissue expressed high levels of CCL28 receptors, particularly CCR3, and this was associated with higher neutrophil accumulation to the gut during STm colitis and to the lung during Ab pneumonia.

### Proinflammatory stimuli and phagocytosis induce expression of CCR3 and CCR10 on neutrophils

We next investigated potential mechanisms underpinning the upregulation of CCR3 and CCR10 in neutrophils. A prior study indicated that a cocktail of proinflammatory cytokines (GM-CSF, IFNγ, TNFɑ) boosted CCR3 expression in human peripheral blood mononuclear cells (PBMCs) from healthy donors (32), and expression of these cytokines is highly induced during STm colitis (**Fig. 1E**) and Ab pneumonia (**Fig. 2J**). We thus stimulated bone marrow neutrophils from wild-type mice (which express low levels of CCR3 and CCR10) with these cytokines, and independently with other pro-inflammatory compounds including lipopolysaccharide (LPS), the protein kinase C activator phorbol 12-myristate 13-acetate (PMA), or the N-formylated, bacterial-derived chemotactic peptide fMLP. Treatment of neutrophils with the GM-CSF + IFNγ + TNFɑ cytokine combination or with LPS induced higher CCR3 (∼40% positive cells) and CCR10 (∼20% positive cells) expression, whereas PMA and fMLP separately yielded more modest yet significant induction (**Fig. 3E, F**).

Phagocytosis of microbes and necrotic debris are critical neutrophil functions at tissue foci of infection and inflammation (33), and gene expression changes are observed in human neutrophils following phagocytosis (34). We thus tested whether CCR3 and CCR10 were induced by phagocytosis, incubating bone marrow neutrophils with latex beads, with or without the aforementioned cytokine cocktail. Although phagocytosis of latex beads alone did not significantly induce neutrophil CCR3 receptor expression (∼7% of neutrophils), latex beads augmented with cytokine cocktail induced CCR3 expression (∼53% of neutrophils vs. ∼30% with cocktail alone; **Fig. 3G**). This synergistic effect of phagocytosis was not noted for CCR10 (**Fig. 3H**).

To further probe the role of phagocytosis in CCR3 expression, we incubated bone marrow neutrophils with live STm for 1h. We found that STm rapidly induced the expression of CCR3 on the neutrophil surface (∼75% of cells; **Fig. 3I**), whereas CCR10 was only minimally induced (**Fig. 3J**). Receptor induction was largely blocked by cytochalasin D, a potent inhibitor of the actin polymerization required for phagocytic uptake (**Fig. 3I, J**). Proinflammatory stimuli and phagocytosis thus enhance the expression of CCR3 and, to a lesser extent, CCR10, on the neutrophil surface.

### CCR3 is stored intracellularly in neutrophils

Neutrophil intracellular compartments and granules harbor enzymes, cytokines, and receptors that are required for rapid responses to pathogens. For example, activation of human neutrophils induces a rapid translocation of complement receptor type 1 (CR1) from an intracellular compartment to the cell surface, increasing its surface expression up to 10-fold (35). As we detected a rapid (within 1h) increase of neutrophil CCR3 surface expression upon STm infection, we hypothesized that CCR3, akin to CR1, may be stored intracellularly in neutrophils, consistent with reports of intracellular CCR3 in eosinophils (36). We found that uninfected bone marrow neutrophils express relatively low surface levels of CCR3 (**Fig. 4A**), but when permeabilized for intracellular staining, most (∼83%) were CCR3⁺, indicating that CCR3 is stored intracellularly (**Fig. 4B**). Upon STm infection *in vitro*, bone marrow neutrophils rapidly increased CCR3 surface expression (up to ∼75%; **Fig. 4A**), consistent with a mobilization of pre-formed receptor from an intracellular compartment (**Fig. 4B**). Intracellular stores of CCR10 were also detected in bone marrow neutrophils under homeostatic conditions, with a small but significant increase during STm infection (**Suppl. Fig. 3E**). However, CCR10 was expressed on the surface of only ∼1% uninfected bone marrow neutrophils, which increased to ∼3% during STm infection (**Suppl. Fig. 3E**). *In vitro*, Ab infection induced higher surface expression of CCR3 on neutrophils (∼26%; **Fig. 4C**), whereas CCR10 did not significantly increase (**Suppl. Fig. 3F**). Most bone marrow neutrophils also expressed intracellular CCR3 (**Fig. 4D**) and CCR10 (**Suppl. Fig. 3F**) during Ab infection. Similar to the *in vitro* findings, neutrophils isolated from bone marrow, blood, and gut tissue of mice orally infected with STm, as well as from bone marrow, blood, and BAL fluid of mice infected with Ab, harbored both intracellular and surface CCR3, albeit at differing levels (**Fig. 4E, F**). Of note, CCR3 surface expression levels were higher on neutrophils isolated from the gut and the BAL relative to other sites (**Fig. 4E, F**). We conclude that CCR3 is stored intracellularly in neutrophils and quickly mobilized to the cell surface upon infection, phagocytosis, and/or cytokine stimulation.

**Figure 4.**
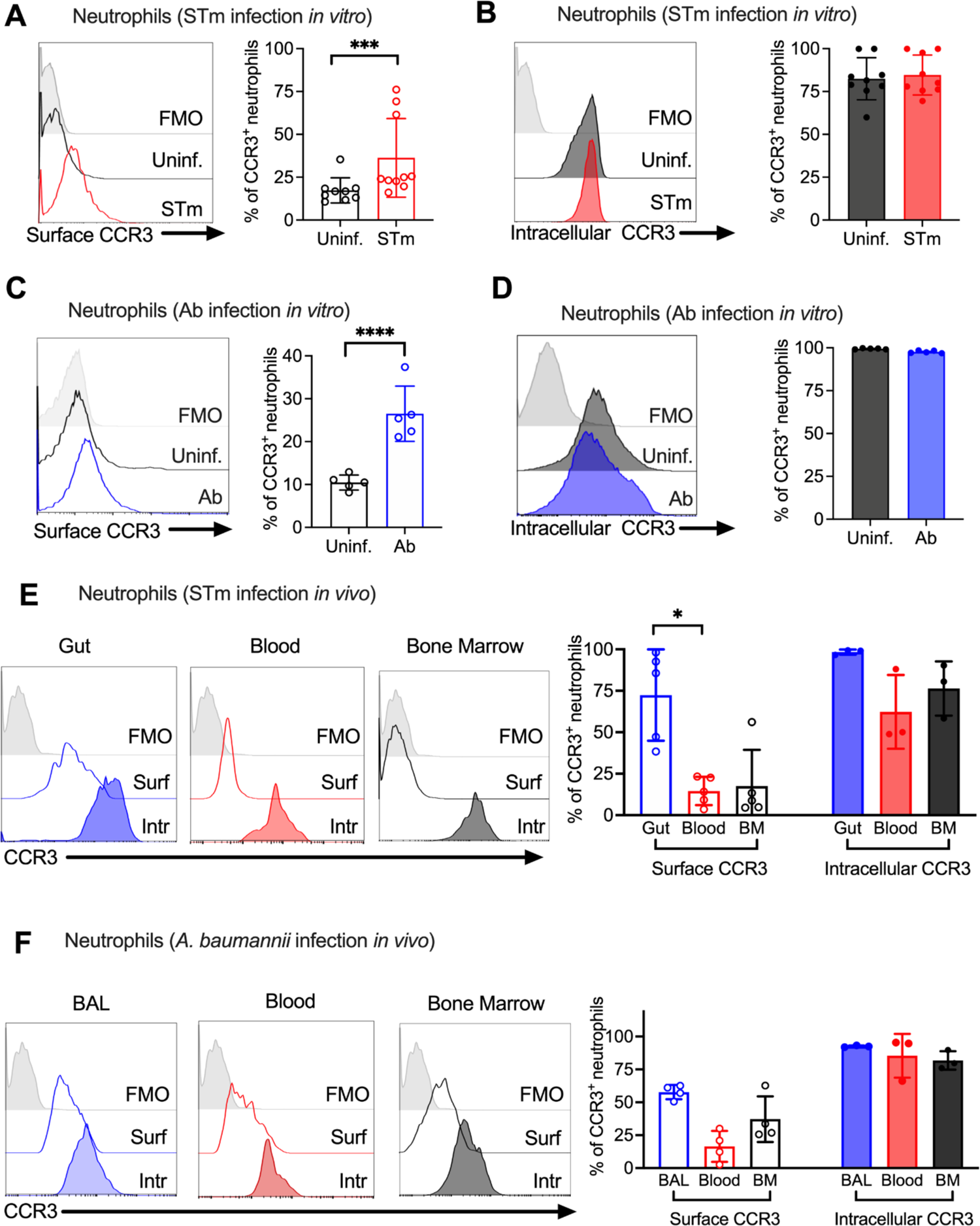
Neutrophil CCR3 is stored in intracellular compartments and rapidly mobilizes to the cell surface during infection. (**A-D**) Bone marrow neutrophils were infected at MOI=10 with (**A, B**) opsonized *Salmonella enterica* serovar Typhimurium (STm) for 1h or (**C, D**) *Acinetobacter baumannii* (Ab) for 4h. (**A, C**) Surface CCR3 or (**B, D**) intracellular CCR3 were detected by flow cytometry. (**E, F**) Neutrophils were obtained from (**E**) the gut, blood, and bone marrow 72h post-infection with STm or (**F**) BAL, blood, and bone marrow 24h post-infection with Ab. Surface (clear histograms) or intracellular (solid histograms) CCR3 expression was analyzed by flow cytometry. (**A-F**) Left panels show representative histograms, and right panels show the percentage of neutrophils expressing CCR3 on their surface (clear bars) or intracellularly (solid bars). Bars represent the mean ± SD. Data was analyzed by paired *t* test (A-D) or one-way ANOVA followed by Dunnett’s multiple comparison test (E and F) on log-transformed data. Significant changes are indicated by \**p* ≤ 0.05, \*\*\**p* ≤ 0.001, \*\*\*\**p* ≤ 0.0001

### CCL28 enhances neutrophil antimicrobial activity, ROS production and NET formation via CCR3 stimulation

Chemokines are essential for neutrophil migration to sites of infection and may regulate additional neutrophil bactericidal effector functions including the production of reactive oxygen species (ROS) and formation of neutrophil extracellular traps (NETs) (37). We tested if CCL28 has chemotactic and/or immunostimulatory activity towards bone marrow neutrophils *in vitro* after boosting their CCR3 surface expression with the cytokine cocktail (GM-CSF + IFNγ + TNFɑ) shown in **Fig. 3**. We incubated the neutrophils either with CCL28, or with the well-known neutrophil chemoattractant CXCL1, or with CCL11/eotaxin, a chemokine that binds CCR3 and is induced in the asthmatic lung to promote eosinophil recruitment (38–40). We found CCL28 promoted neutrophil chemotaxis, though not as potently as CXCL1, while CCL11 had no significant effect (**Fig. 5A**).

**Figure 5.**
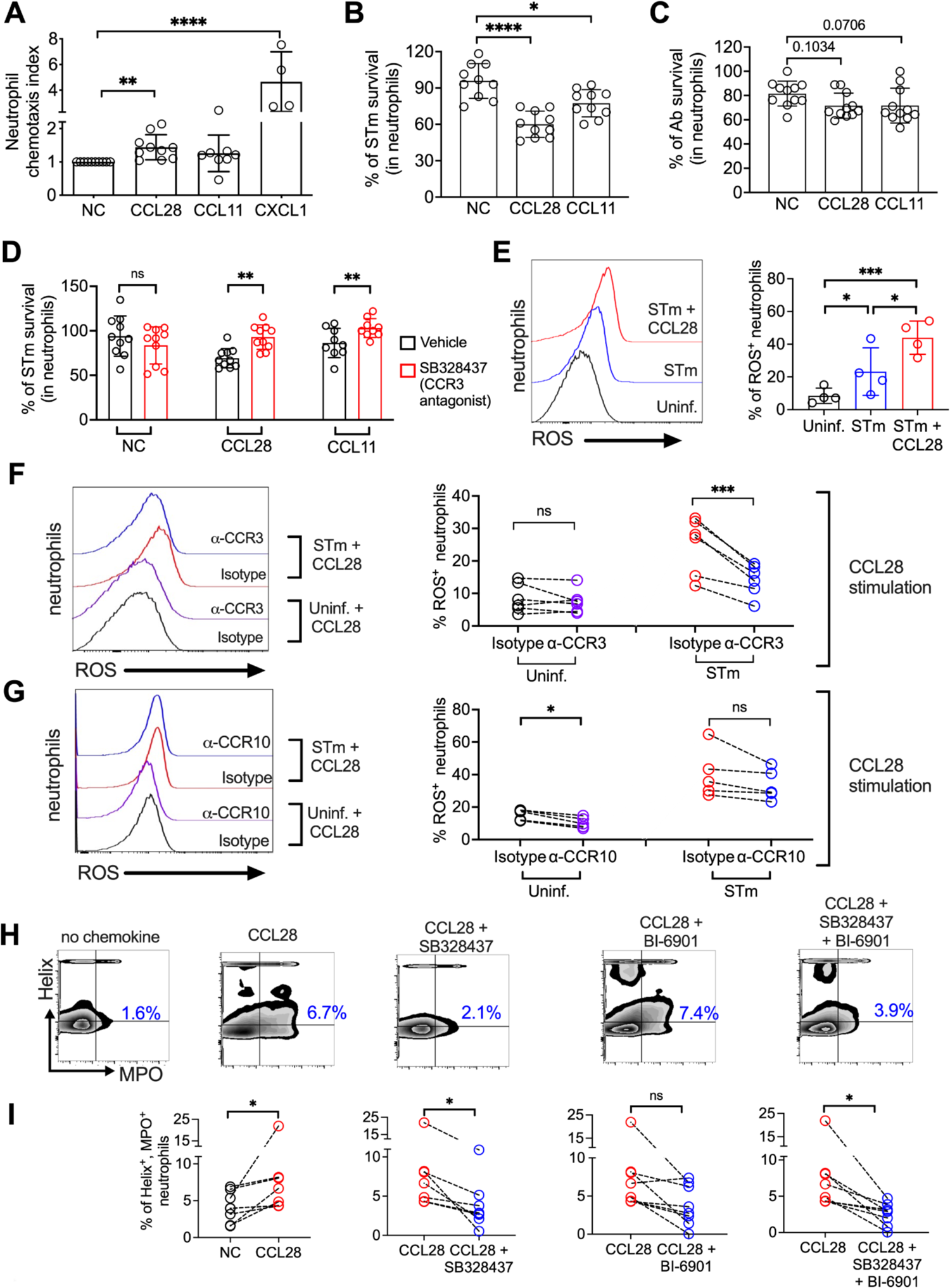
CCL28 enhances neutrophil antimicrobial activity. (**A**) Murine bone marrow neutrophils were stimulated with IFNγ + TNFɑ + GM-CSF for 4h before adding 1×10^6^ cells to the upper compartment of a transwell chamber for chemotaxis assays. Each of the chemokines (CCL28, CCL11, or CXCL1), or no chemokine (NC), were placed in separate lower compartments. The transwell plate was incubated for 2h at 37°C. Cells that migrated to the lower compartment were enumerated by flow cytometry. Neutrophil chemotaxis index was calculated by taking the number of cells that migrated in response to a chemokine and dividing it by the number of cells that migrated in the absence of a chemokine. Data are from four independent experiments. (**B, C**) Infection of bone marrow neutrophils. (**B**) Opsonized STm (1×10^7^ CFU) or (**C**) opsonized Ab (1×10^7^ CFU) were cultured alone, or added to bone marrow neutrophils (1×10^6^ cells) stimulated with CCL28, CCL11, or no chemokine, for 2.5h (STm) or 4.5h (Ab) at 37°C. CFU were enumerated by plating serial dilutions. Percentage of bacterial survival was calculated for each condition by taking the CFU from bacteria incubated with neutrophils and dividing it by the CFU from bacteria incubated without neutrophils, multiplied by 100. Data shown for each infection comprise three independent experiments. Bars represent the mean ± SD. (**D**) The effect of the CCR3 antagonist SB328437 on neutrophil-mediated STm killing was evaluated by performing the experiment as described in panel (**B**), with or without the antagonist. Data shown comprise three independent experiments. (**E-G**) ROS production (H_2_DCFDA conversion to fluorescent DCF) detected by flow cytometry in bone marrow neutrophils infected with STm as described in panel (**B**). In (**F, G**), cells were stimulated with CCL28 in the presence of an anti-CCR3 antibody, an anti-CCR10 antibody, or isotype controls. Left panels show representative histograms, and right panels show the percentage of ROS^+^ neutrophils in the indicated treatment groups. (**H, I**) NET formation (Helix^+^ MPO^+^ neutrophils) detected by flow cytometry in human neutrophils activated with platelets. As indicated, cells were unstimulated (NC), stimulated with CCL28 alone, or with CCL28 and the CCR3 antagonist SB328437 and/or the CCR10 antagonist BI-6901. (**H**) Representative contour plots, and (**I**) percentage of Helix^+^ MPO^+^ neutrophils in the indicated treatment groups. (**A**-**E**) Bars represent the mean ± SD. (**A**) Log-transformed data were analyzed by non-parametric ANOVA (Kruskal-Wallis’s test), assuming non-equal SD given the differences in the variance among the groups, followed by Dunn’s multiple comparisons test. (**B, C**) Log-transformed data were analyzed by one-way ANOVA for paired samples, applying the Greenhouse-Geisser correction given the differences in the variance among the groups. Dunnett’s multiple comparison test was performed to compare the treatment groups only with the control group. (**D, I**) Log-transformed data were analyzed by paired *t* test. (**E-G**) Log-transformed data were analyzed by one-way ANOVA for paired samples. Greenhouse-Geisser correction was applied in **F** and **G** given the differences in variance among the groups. Tukey’s multiple comparison test was performed to compare all conditions to each other. Significant changes are indicated by \**p* ≤ 0.05, \*\**p* ≤ 0.01, \*\*\**p* ≤ 0.001 \*\*\*\**p* ≤ 0.0001. ns = not significant.

To test whether CCL28 stimulation enhanced neutrophil effector function, we incubated STm with bone marrow neutrophils for 2.5h with or without CCL28 (50nM) or CCL11 (50nM), then quantified bacterial killing. Stimulation with CCL28 strongly increased neutrophil bactericidal activity against STm, with clearance of ∼40% of the bacterial inoculum, compared to only ∼10% clearance seen with unstimulated neutrophils (**Fig. 5B**). Neutrophils stimulated with CCL11 displayed an intermediate phenotype (∼25% bacterial killing). As expected, neither chemokine exhibited direct antimicrobial activity against STm (**Suppl. Fig. 1D**). In stark contrast to findings with STm, *ex vivo* neutrophil killing of Ab was not significantly enhanced by CCL28 or CCL11 treatment (**Fig. 5C**). Thus, even though CCL28 modulates neutrophil accumulation in the lung during Ab infection (**Fig. 2D-J**), a plausible contributing factor to why CCL28 fails to reduce pathogen burden in the lung (**Fig. 2B**) is that CCL28 stimulation does not enhance neutrophil bactericidal activity against Ab.

Our data indicate that CCR3 is the CCL28 receptor that is most highly expressed in neutrophils during STm infection (**Fig. 3I and Fig. 4**). We next tested whether the CCL28-mediated increase in neutrophil bactericidal activity could be reversed in the presence of the small molecule SB328437, a CCR3 antagonist (41). SB328437 reversed the effect of both CCL28 and CCL11 on neutrophils, corroborating receptor specificity (**Fig. 5D**). An important mechanism of bacterial killing is the production of ROS (42), a process that is triggered in response to infection and enhanced by proinflammatory stimuli including cytokines and chemokines (43). We measured ROS production by incubating neutrophils with the cell-permeable probe 2’,7’-dichlorodihydrofluorescein diacetate (H_2_DCFDA), and found that CCL28 stimulation enhanced neutrophil ROS production during STm infection (**Fig. 5E**). Importantly, the increased ROS production triggered by CCL28 was reversed when neutrophils were incubated with an anti-CCR3 blocking antibody (**Fig. 5F**), but not with an anti-CCR10 blocking antibody (**Fig. 5G**).

In addition to their direct antimicrobial activity, ROS trigger other neutrophil responses, including NET formation (43). NETs can be induced by a variety of stimuli including microbial products, inflammatory cytokines and chemokines, immune complexes and activated platelets (44). To determine whether CCL28 enhances NET formation, we incubated human neutrophils with activated platelets with or without CCL28 and evaluated the formation of DNA-MPO complexes by flow cytometry. Incubation of human neutrophils with activated platelets and CCL28 increased the percentage of DNA-MPO complexes compared to neutrophils not stimulated with CCL28 (**Fig 5H, 5I**). The effect of CCL28 on platelet-activated NET formation was largely mediated by CCR3, as incubation with the CCR3 antagonist SB328437 significantly reduced the percentage of neutrophils with DNA-MPO complexes (**Fig 5H, 5I**). In contrast, incubation with the CCR10 antagonist BI-6901 did not significantly reduce NET formation, and incubation with both CCR3 and CCR10 antagonists exhibited a similar effect as the CCR3 antagonist alone (**Fig 5H, 5I**). Together, these results demonstrate that CCL28 enhances neutrophil ROS production and NET formation primarily in a CCR3-dependent manner.

## Discussion

The mucosal immune response serves to maintain tissue homeostasis and to protect the host against invading pathogens. Here we discovered that the chemokine CCL28 contributes to neutrophil accumulation and activation in the mucosa in two infection models: gastrointestinal infection with *Salmonella*, and lung infection with *Acinetobacter*.

Consistent with our initial observation that *Ccl28*^-/-^ mice exhibit higher lethality during STm infection (9), we found higher intestinal colonization and extraintestinal dissemination of STm in *Ccl28*^-/-^ mice than their wild-type littermates (**Fig. 1**). This beneficial role for CCL28 was negligible when the pathogen was inoculated intraperitoneally to bypass the gut mucosa (**Suppl. Fig. 1**). Although it has been reported that CCL28 exerts direct antimicrobial activity against some bacteria and fungi (18), the chemokine does not directly inhibit STm wild-type *in vitro* (**Suppl. Fig. 1**). Moreover, even though the CCL28 receptors CCR3 and CCR10 are expressed on eosinophils and on B and T cells (8, 31, 45), none of these cell types appeared to be responsible for the observed protective role of CCL28 during *Salmonella* infection. Eosinophils are not a major component of the host response to *Salmonella*, and we observed comparable numbers of B and T cells in the gut during homeostasis, as well as during *Salmonella* infection, in wild-type and *Ccl28*^-/-^ mice (**Suppl. Fig. 2**).

Neutrophils are a hallmark of inflammatory diarrhea. In the *Salmonella* colitis model, neutrophils are rapidly recruited to the gut following infection. We found that neutrophil numbers were significantly reduced in the mucosa of infected *Ccl28*^-/-^ mice relative to wild-type mice (**Fig. 1**), thereby identifying CCL28 as a key factor for neutrophil accumulation during infection. Neutrophils migrate from the bone marrow to the blood to infected sites following a chemokine gradient (37), and express chemokine receptors CXCR1, CXCR2, CXCR4 and CCR2, as well as CCR1 and CCR6 under certain circumstances (46). CXCR2 is a promiscuous receptor that binds to the chemokines CXCL1, 2, 3, 5, 6, 7, and 8 (47), whereas CXCR1 only binds CXCL6 and CXCL8 (37). Activation of CXCR2 induces mobilization of neutrophils from the bone marrow to the bloodstream and participates in the release of NETs (48). Engagement of CXCR1 and CXCR2 also triggers signaling pathways resulting in the production of cytokines and chemokines that amplify neutrophil responses (25). Following extravasation to the site of infection, neutrophils downregulate CXCR2 and upregulate CCR1, 2, and 5, which cumulatively boosts neutrophil ROS production and phagocytic activity (37). Our results indicate that CCL28 contributes to neutrophil accumulation and activation (**Fig. 1**), and that its receptors CCR3 and CCR10 are upregulated in the mucosa during infection, where up to ∼75% of neutrophils express surface CCR3 (**Fig. 3**). The reciprocal regulation of CXCR2 and CCR3/CCR10 in neutrophils and each receptor’s contribution to neutrophil migration and retention during infectious colitis need to be elucidated by further studies.

Although an initial study concluded CCR3 was absent on neutrophils (31), two subsequent studies reported CCR3 expression on human neutrophils isolated from the lung of patients with chronic lung disease (32) and on neutrophils isolated from the BAL fluid of mice infected with influenza (49). CCR10 expression in neutrophils has not been previously reported. Our study demonstrates that a substantial number of neutrophils isolated from infected mucosal sites express CCR3 as well as CCR10 on their surface, whereas resting neutrophils do not express these receptors on their surface (**Fig. 3**). As we detected CCR3 on the neutrophil surface quite rapidly after infection, we predicted that the receptor was stored in intracellular compartments, akin to what was found in eosinophils (36). Indeed, neutrophils isolated from bone marrow, blood, and infected mucosal tissue were all positive for CCR3 intracellular staining (**Fig. 4**), and we could recapitulate the increase of surface receptor expression *in vitro* by incubating bone marrow neutrophils with proinflammatory stimuli (LPS, or the cytokines GM-CSF + IFNγ + TNFɑ), or following phagocytosis of bacterial pathogens (**Fig. 3**). In all cases, upregulation of surface CCR3 on neutrophils was more robust than that of CCR10. CCL28 stimulation of bone marrow neutrophils *in vitro* increased their antimicrobial activity and ROS production when infected with *Salmonella*, which was reverted by blocking CCR3 but not CCR10 (**Fig. 5**). Platelet-activated neutrophils stimulated with CCL28 also showed enhanced NET formation, again largely in a CCR3-dependent fashion (**Fig. 5**). Thus, CCL28 is a potent activator of neutrophils, and it appears to do so primarily via CCR3. Further studies with receptor knock-out mice are needed to determine the contribution of each CCL28 receptor to the *in vivo* phenotypes.

A reduction of neutrophil accumulation was also observed in the BAL and lung of *Ccl28*^-/-^ mice during *Acinetobacter* infection (**Fig. 2**), and neutrophils recruited to the lung harbored surface CCR3 and CCR10 (**Fig. 3, 4**). However, the functional consequences of CCL28 deficiency was strikingly different in this model, as *Ccl28*^-/-^ mice were protected during Ab pneumonia. Most *Ccl28^-/-^* mice survived until the experiment’s arbitrary endpoint at Day 10 post-infection, whereas nearly all wild-type littermates succumbed by Day 2 (**Fig. 2**). The lung, possessing a thin, single-cell alveolar layer to promote gas exchange, is less resilient than the intestine to neutrophil inflammation before losing barrier integrity and critical functions. Thus, even though insufficient neutrophil recruitment can lead to life-threatening infection, extreme accumulation of neutrophils can result in excessive inflammatory lung injury (50). The high survival of *Ccl28^-/-^* mice infected with Ab indicates that CCL28 may be detrimental for the host at least in the context of some pulmonary infection. While functioning neutrophils have been described to play a role in controlling *Acinetobacter* infection (29, 51, 52), an exaggeration of neutrophil recruitment to the *Acinetobacter*-infected lung is deleterious (53–55). For example, in one relevant *Acinetobacter* pneumonia study, depletion of alveolar macrophages increased neutrophil infiltration, promoted extensive tissue damage, and increased systemic dissemination of *Acinetobacter* (56). Of note, in contrast to *Salmonella*, stimulation with CCL28 did not enhance neutrophil antimicrobial activity against *Acinetobacter*, which may partly explain the lack of a CCL28-mediated protective response (**Fig. 5**). The bacteriological mechanism(s) by which *Acinetobacter* may be resistant to CCL28-dependent neutrophil antimicrobial responses require further investigation.

Even though CCL28 exhibited direct antimicrobial activity against *Acinetobacter*, higher concentrations of CCL28 (1μm) are needed for protection, and even so are not sufficient against higher pathogen burden (**Suppl. Fig. 1**). Our findings are consistent with prior studies showing that the impact of the host response to infection can be context dependent, and that some immune components mediate different outcomes in the gut and in the lung. For example, *Cxcr2*^-/-^ mice exhibit a defect in neutrophil recruitment that is detrimental during *Salmonella* infection (57), but protective during lung infection with *Mycobacterium tuberculosis*, due to reduced neutrophil recruitment and reduced pulmonary inflammation (58). Similarly, CCL28-dependent modulation of neutrophil accumulation and activation during infection can be protective or detrimental depending on the pathogen and on the site of infection.

Altogether, this study demonstrates that CCL28 plays an important role in the mucosal response to pathogens through neutrophil accumulation to the site of infection, that neutrophils isolated from the infected mucosa express the CCL28 receptors CCR3 and CCR10, and that CCL28 enhances neutrophil activation, ROS production and NET formation, primarily through CCR3. These findings could have implications for other infectious and non-infectious diseases where neutrophil recruitment plays a major role, and may lead to the identification of new CCL28-targeted therapies to modulate neutrophil function and mitigate collateral damage.

## Materials and methods

### Generation and breeding of *Ccl28*^-/-^ mice

The first colony of *Ccl28*^-/-^ mice was described in a prior manuscript (9) and used for initial studies at UC Irvine. At UC San Diego, we generated a new colony of *Ccl28*^-/-^ mice with Cyagen Biosciences (Santa Clara, California), using CRISPR/CAS9 technology. Exons 1 and 3 were selected as target sites, and two pairs of gRNA targeting vectors were constructed and confirmed by sequencing. The gRNA and Cas9 mRNA were generated by *in vitro* transcription, then co-injected into fertilized eggs for knockout mouse production. The resulting pups (F0 founders) were genotyped by PCR and confirmed by sequencing. F0 founders were bred to wild-type mice to test germline transmission and for F1 animal generation. Tail genotyping of offspring was performed using the following primers: F: 5’-TCATATACAGCACCTCACTCTTGCCC-3’, R: 5’-GCCTCTCAAAGTCATGCCAGAGTC-3’ and He/Wt-R: 5’-TCCCGGCCTTGAGTATGTTAGGC-3’. The expected product size for the wild-type allele is 466 bp and for the knockout allele is 700 bp.

All mouse experiments were conducted with cohoused wild-type and *Ccl28*^-/-^ littermate mice, and were reviewed and approved by the Institutional Animal Care and Use Committees at UC Irvine and UC San Diego.

### *Salmonella* infection models

For the *Salmonella* colitis model, 8-10 week-old male and female mice were orally gavaged with 20mg streptomycin 24h prior to oral gavage with 10^9^ colony-forming units (CFU) of *Salmonella enterica* serovar Typhimurium strain IR715 (a fully virulent, nalidixic acid-resistant derivative of ATCC 14028s) (59), as previously described (17, 60). Mice were euthanized at 72h post-infection, then colon content, spleen, mesenteric lymph nodes, Peyer’s patches, and bone marrow were collected, weighed, homogenized, serially diluted, and plated on Miller Lysogeny broth (LB) + Nal (50µg/mL) agar plates to enumerate *Salmonella* CFU. For the *Salmonella* bacteremia model, mice were injected intraperitoneally with 10^3^ CFU. Mice were euthanized at 72h post-infection, then blood, spleen, and liver were collected to determine bacterial counts.

### *Acinetobacter* infection model

For the murine pneumonia model, *Acinetobacter baumannii* strain AB5075 was cultured in Cation-Adjusted Mueller-Hinton Broth (CA-MHB) overnight, then subcultured the next day to an OD_600_ of ∼0.4 (1×10^8^ CFU/mL; mid-logarithmic phase). These cultures were centrifuged at 3202x*g*, the supernatant was removed, and pellets were resuspended and washed in an equal volume of 1x Dulbecco’s PBS (DPBS) three times. The final pellet was resuspended in 1x DPBS to yield a suspension of 2.5 x 10^9^ CFU/mL. Using an operating otoscope (Welch Allyn), mice under 100 mg/kg ketamine (Koetis) + 10 mg/kg xylazine (VetOne) anesthesia were inoculated intratracheally with 40 μL of the bacterial suspension (10^8^ CFU/mouse). Post-infection mice were recovered on a sloped heating pad. For analysis of bacterial CFU, mice were sacrificed at 24h post-infection, the BAL was collected, and serial dilutions were plated on LB agar to enumerate bacteria (26).

### CCL28 ELISA

Fresh fecal samples were collected at 96h post-infection from wild-type mice. Fecal pellets were weighed, resuspended in 1 mL of sterile PBS containing a protease inhibitor cocktail (Roche), and incubated at room temperature shaking for 30 min. Samples were centrifuged at 9391 x *g* for 10 min, supernatants were collected, then analyzed to quantify CCL28 using a sandwich ELISA kit (BioLegend).

### Cell extraction and analysis

For the *Salmonella* colitis model, the terminal ileum, cecum, and colon were collected at 48h post-infection. All tissues were kept in IMDM medium supplemented with 10% FBS and 1% antibiotic/antimycotic. Next, the Peyer’s patches were removed, and the intestinal fragments were cut open longitudinally and washed with HBSS supplemented with 15 mM HEPES and 1% antibiotic/antimycotic. Then, the tissue was shaken in 10 mL of an HBSS/ 15 mM HEPES/ 5 mM EDTA/ 10% FBS solution at 37 °C in a water bath for 15 min. The supernatant was removed and kept on ice. The remaining tissue was cut into small pieces and digested in a 10 mL mixture of collagenase (Type VII, 1 mg/mL), Liberase (20 µg/mL), and DNAse (0.25 mg/mL) in IMDM medium for 15 min in a shaking water bath at 37 °C. Afterwards, the supernatant and tissue fractions were strained through a 70 µm cell strainer and pooled, and the extracted cells were used for flow cytometry staining. For the *A. baumannii* infection model, the lungs were collected and processed as described for the gut. BAL was collected, centrifuged, and pellets were washed with 1x PBS. The obtained cells were used for flow cytometry staining. Briefly, cells were blocked with a CD16/32 antibody (Bio-Legend), stained with the viability dye eFluor780 (Thermo Fisher), then extracellularly stained using the following monoclonal antibodies: anti-mouse CD45 (clone 30-F11), anti-mouse CD3 (clone 17A2), anti-mouse CD4 (clone RM4-5), anti-mouse CD8α (clone 53-6.7), anti-mouse CD19 (clone 1D3/CD19), anti-mouse Ly6G (clone 1A8), anti-mouse CD11b (clone M1/70) from BioLegend; anti-mouse CCR3 (clone FAB729P) and anti-mouse CCR10 (clone FAB2815A) from R&D Systems. After staining, cells were fixed for 20 min with 4% paraformaldehyde (Fixation buffer; BioLegend). When intracellular staining was performed, cells were permeabilized (Permeabilization buffer; BioLegend), and the staining was performed in the same buffer following the manufacturer’s instructions. Cells were analyzed on an LSRII flow cytometer (BD Biosciences) and the collected data were analyzed with FlowJo v10 software (TreeStar). For analysis of human neutrophils, whole-blood samples were collected in ethylenediaminetetraacetic acid (EDTA) for cellular analyses. Whole blood cell staining was performed using an Fc receptor blocking solution (Human TruStain FcX; BioLegend), the viability dye eFluor780 (Thermo Fisher), and the following conjugated monoclonal antibodies: PerCP/Cy5.5 anti-human CD45 antibody (clone HI30), Pacific Blue anti-mouse/human CD11b antibody (clone M1/70), FITC anti-human CD62L antibody (clone DREG-56), from BioLegend; PE anti-human CCR3 antibody (clone 61828), and APC anti-human CCR10 antibody (clone 314305) from R&D Systems. Samples were analyzed by flow cytometry using an LSR Fortessa flow cytometer (BD Biosciences), and data was analyzed using FlowJo v10 software.

### *In vitro* neutrophil stimulation

Neutrophils were obtained from the bone marrow of C57BL/6 wild-type mice using the EasySep Mouse Enrichment Kit (STEMCELL), following the manufacturer’s instructions. After enrichment, 1×10^6^ neutrophils were seeded in a round-bottom 96-well plate with RPMI media supplemented with 10% FBS, 1% antibiotic/antimycotic mix, and HEPES (1mM) (Invitrogen). For stimulation, cells were incubated with the following concentrations of recombinant mouse cytokines: TNFɑ (100 ng/mL), IFNγ (500 U/mL) and GM-CSF (10 ng/mL), all from R&D systems. Recombinant cytokines were added alone or in combination. For indicated experiments, polystyrene beads (Thermo Fisher) were added to neutrophils at a 1:1 ratio. Cells were incubated for 4 hours at 37°C and 5% CO_2_. After incubation, cells were recovered by centrifugation, washed with PBS, and processed for flow cytometry as described above.

### Chemotaxis assay

Enriched neutrophils from the bone marrow of wild-type mice were stimulated with a cocktail of mouse recombinant cytokines (TNFɑ, IFNγ, GM-CSF), as described above, to induce receptor expression. After stimulation, cells were washed twice with PBS, resuspended in RPMI media supplemented with 0.1% BSA (w/v) to a final concentration of 1×10^7^ cells/mL, and 100 μL of the cell suspension were placed in the upper compartment of a Transwell chamber (3.0 μm pore size; Corning Costar). 50 nM of recombinant mouse CCL28, CCL11 (R&D Systems), or CXCL1 (Peprotech) were placed into the lower compartment of a Transwell chamber. The Transwell plate was then incubated for 2h at 37 °C. The number of cells that migrated to the lower compartment was determined by flow cytometry. The neutrophil chemotaxis index was calculated by dividing the number of cells that migrated in the presence of a chemokine by the number of cells that migrated in control chambers without chemokine stimulation.

### Neutrophil *in vitro* infection and bacterial killing assays

Bone marrow neutrophils were obtained from mice as described above. *S.* Typhimurium and *A. baumannii* were grown as described in the respective mouse experiment sections. For STm infection, bacteria were then opsonized with 20% normal mouse serum for 30 min at 37 °C. After neutrophils were enriched, 1×10^6^ neutrophils were seeded in a round-bottom 96-well plate with RPMI media supplemented with FBS (10%), and bacteria were added at a multiplicity of infection (MOI)= 10. The plate was centrifuged to ensure interaction between cells and bacteria. After 30 min of contact with STm, the media was changed and RPMI media supplemented with FBS (10%) and gentamicin (100 μg/mL) was added, and cells were incubated for 30 min more. For Ab infection, 1×10^6^ neutrophils were seeded in a round-bottom 96-well plate with RPMI media supplemented with FBS (10%), and bacteria were added at a multiplicity of infection (MOI)= 10. The plate was centrifuged to ensure interaction between cells and bacteria, and cells were incubated for 4 h at 37 °C and 5% CO2. For analysis of CCR3 and CCR10 expression, cells were recovered by centrifugation, washed with PBS, and processed for flow cytometry as described above. For inhibition of phagocytosis, bone marrow neutrophils were pre-incubated with cytochalasin D (10 µM) in DMSO (0.1%), or DMSO (vehicle), for 30 min prior to infection with opsonized *S*. Typhimurium for 1h at an MOI=10. For killing assays, recombinant mouse CCL28 (50nM) (45) and CCL11 (25nM) (61) (R&D systems) were added to neutrophils. When indicated, the CCR3 receptor antagonist SB328437 (Tocris Bioscience) was added at a final concentration of 10 μM (41). Neutrophils infected with STm were incubated for 2.5h and neutrophils infected with *A. baumannii* were incubated for 4.5h at 37 °C and 5% CO_2_. After incubation, 1:2 dilution was performed with PBS supplemented with 2% Triton X-100 and then serial dilution was performed and plated on LB agar to enumerate bacteria. To calculate the percentage of bacterial survival, the number of bacteria recovered in the presence of neutrophils was divided by the number of bacteria recovered from wells that contained no neutrophils, then multiplied by 100.

### Reactive oxygen species (ROS) production

Neutrophils were obtained from the bone marrow of C57BL/6 wild-type mice using the EasySep Mouse Enrichment Kit (STEMCELL Technologies), following the manufacturer’s instructions. After enrichment, 2.5 x 10^6^ cells/mL were resuspended in phenol red-free RPMI media (Gibco) supplemented with 10% FBS (Gibco), and HEPES (1mM, Invitrogen). The cells were incubated in presence of 2’,7’-dichlorodihydrofluorescein diacetate (H_2_DCFDA, 25 mM) (Invitrogen) for 30min at 37 °C and 5% of CO_2_, as previously described (62). After incubation with H_2_DCFDA, neutrophils were infected with STm as described above, then incubated for 4h with mouse recombinant CCL28 (50 nM), anti-mouse CCR3 antibody (5 mg/1×10^6^ cells, clone 83103), anti-mouse CCR10 antibody (5 mg/1×10^6^ cells, clone 248918) or anti-rat IgG2A (5 mg/1×10^6^ cells, clone 54447), all from R&D Systems. Neutrophils were analyzed by flow cytometry to determine ROS production using a BD FACSCanto flow cytometer, and data was analyzed using the FlowJo v10 software.

### Neutrophil extracellular traps (NETs) production

Whole-blood samples were collected from healthy donors recruited at a tertiary care center in Mexico City (Instituto Nacional de Ciencias Médicas y Nutrición Salvador Zubirán). Healthy donors signed an informed consent form before inclusion in the study, and the protocol was approved by the Instituto Nacional de Ciencias Médicas y Nutrición Salvador Zubirán ethics and research committees (Ref. 3341) in compliance with the Helsinki declaration. Neutrophils were obtained from peripheral blood of healthy voluntary donors using the EasySep Direct Human Neutrophil Isolation Kit (STEMCELL Technologies), following the manufacturer’s instructions. In parallel, platelets from human peripheral blood were isolated as described (63). Briefly, whole blood was centrifuged at 200*g* for 10 minutes at 4 °C, and plasma was recovered and then centrifuged again at 1550*g* for 10 minutes at 4 °C. The cell pellet was resuspended in RPMI media supplemented with 10% FBS (4 x 10^7^ cells/mL) and then incubated with LPS (5 mg/mL) for 30 minutes at 37 °C to induce platelet activation (64). Subsequently, neutrophils were incubated with autologous activated platelets (1:10 ratio) (65) for 2.5h and human recombinant CCL28 (50 nM) (BioLegend), the CCR3 antagonist SB328437 (10 mM, Tocris Bioscience) and/or the CCR10 antagonist BI-6901 (20 mM, Boehringer-Ingelheim) were added as indicated. Cells were then incubated with the DNA-binding dye Helix-NP Green (10 nM, BioLegend), human anti-myeloperoxidase (MPO)-Biotin antibody (clone MPO421-8B2, Novus Biologicals), and APC/Cy7 streptavidin (BioLegend). Samples were analyzed using an LSR Fortessa flow cytometer (BD Biosciences) to determine the presence of DNA-MPO complexes (66), and data were analyzed using FlowJo v10 software.

### Growth of bacteria in media supplemented with recombinant chemokines

*S.* Typhimurium wild-type, *S.* Typhimurium *phoQ* mutant, and *Escherichia coli* K12 were grown in LB broth overnight at 37 °C. *Acinetobacter baumannii* was cultured in Cation-Adjusted Mueller-Hinton Broth (CA-MHB) under the same conditions. The following day, cultures were diluted 1:100 in LB and grown at 37 °C for 3 hr, subsequently diluted to ∼0.5 x 10^6^ CFU/mL or ∼0.5 x 10^9^ CFU/mL in 1 mM potassium phosphate buffer (pH 7.2), then incubated at 37 °C in the presence or absence of recombinant murine CCL28 (BioLegend) at the indicated concentrations. After 2h, samples were plated onto LB agar to enumerate viable bacteria. In other assays, *S.* Typhimurium was grown as described above and ∼1×10^7^ CFU/mL were incubated at 37 °C for 2.5h in the presence or absence of recombinant murine CCL28 (50 nM) (45) or CCL11 (25 nM) (61) in RPMI medium supplemented with 10% FBS. After incubation, samples were plated onto LB + Nal agar to enumerate viable bacteria.

### RNA extraction and qPCR

Total RNA was extracted from mouse cecal or lung tissue using Tri-Reagent (Molecular Research Center). Reverse transcription of 1 μg of total RNA was performed using the SuperScript VILO cDNA Synthesis kit (Thermo Fisher Scientific). Quantitative real-time PCR (qRT-PCR) for the expression of *Actb* (β-actin), *Cxcl1, Tnfa, Ifng, Csf2, Il1b,* and *Il17a* was performed using the PowerUp SYBR Green Master Mix (Applied Biosystems) on a QuantStudio 5 Real-Time PCR System (Thermo Fisher Scientific). Gene expression was normalized to *Actb* (β-actin). Fold changes in gene expression were relative to uninfected controls and calculated using the *ΔΔ*Ct method.

### Histopathology

Cecal and lung tissue samples were fixed in 10% buffered formalin, embedded in paraffin according to standard procedures, and sectioned at 5 μm. Pathology scores of cecal and lung samples were determined by blinded examinations of hematoxylin and eosin (H&E)-stained sections. Each cecal section was evaluated using a semiquantitative score as described previously (67). Lung inflammation was assessed by a multiparametric scoring based on previous work (68).

### Immunofluorescence

Deparaffinized lung sections were stained with a purified rat anti-mouse Ly6G antibody (clone 1A8, BioLegend) according to standard immunohistochemical procedures. Ly6G+ cells were visualized by a goat anti-rat secondary antibody (Invitrogen). Cell nuclei were stained with DAPI in SlowFade Gold Antifade Mountant (Invitrogen). Slides were scanned on a Zeiss Axio Scan.Z1 slide scanner and whole lung scans were evaluated with QuPath analysis software (69). Ly6G+ cells per mouse were quantified by averaging the neutrophil numbers of 3 consecutive high-power fields in regions with moderate to severe inflammation.

### Statistical analysis

Statistical analysis was performed with GraphPad Prism 9. CFU data, percentage of CCR3^+^ or CCR10^+^ neutrophils *in vivo* and *in vitro*, and data from functional neutrophil assays were transformed to Log10 and passed a normal distribution test before running statistical analyses.

CFU from *in vivo* infection experiments, cytokine secretion, qPCR data, and neutrophils numbers were compared by Mann-Whitney U test. Survival curves were compared by the Log-rank (Mantel-Cox) test. Data that were normally distributed were analyzed by one-way ANOVA for independent samples or paired samples, depending on the experimental setup. Dunnett’s multiple comparisons test was applied when we compared the different conditions to a single control group, while Tukey’s multiple comparison test was performed when we compared each condition with each other. Greenhouse-Geisser correction was applied when there were differences in the variance among the groups. Data from chemokine migration were analyzed by a non-parametric ANOVA (Kruskal-Wallis’s test), assuming non-equal SD given the differences in the variance among the groups and followed by Dunn’s multiple comparisons test. Paired *t* test was used when only two paired experimental groups were compared. A p value equal to or below 0.05 was considered statistically significant. * indicates an adjusted p value ≤0.05, ** p value ≤0.01, *** p value ≤0.001, **** p value ≤0.0001.

## Acknowledgements

This work was supported by the NIH (Public Health Service Grants AI121928) to MR, by a pilot project award from the NIAID Mucosal Immunology Studies Team (MIST) to APL, and by a grant from the InnovaUNAM of the National Autonomous University of Mexico (UNAM) and Alianza UCMX of the University of California, to APL and MR. RRG was partly supported by a fellowship from the Max Kade Foundation and by a fellowship from the Crohn’s and Colitis Foundation. MHL was partly supported by NIH training grant T32 DK007202. ND was supported by NIH training grant NIH 5T32HD087978-05 and NIH NIAID grant 1-U01-AI124316. GTW was supported by NIH training grant T32AI007036. KM was partly supported by São Paulo Research Foundation (FAPESP) Grants 2019/14833-0 and 2018/22042-0. RCD, SR-R, and VAS-H were supported by a Grad School Fellowship from CONACyT. MR and VN are supported by Public Health Service Grant AI145325. Work in JLM-M lab was supported by CONACyT-FOSISS grant A3-S-36875 and UNAM-DGAPA-PAPIIT Program grant IN213020. Work in MR lab is also supported by NIH Grants AI126277, AI145325, AI154644, AI096528, and by the Chiba University-UCSD Center for Mucosal Immunology, Allergy, and Vaccines. MR holds an Investigator in the Pathogenesis of Infectious Disease Award from the Burroughs Wellcome Fund. We would like to thank Dr. Albert Zlotnik for his thoughtful suggestions on the project over the years. We also acknowledge Boehringer-Ingelheim Pharma GmbH & Co. KG for the kind gift of the CCR10 antagonist BI-6901, and the Histology Core at the La Jolla Institute for Immunology.

## Author contributions

APL and MR conceived the overall study. APL, SS, MHL, ND, SLB, RRG, KM, GTW, S-PN, VN, JLM-M, and MR designed the experiments and analyzed the data. APL, SS, MHL, ND, SLB, RRG, KM, GTW, VAS-H, RC-D, and SR-R performed experiments. RRG analyzed the histopathology. APL, S-PN, VN, and MR wrote the paper. APL, S-PN, VN, and MR provided supervision and funding support.

## SUPPLEMENTAL FIGURES

**Supplementary Figure 1.**
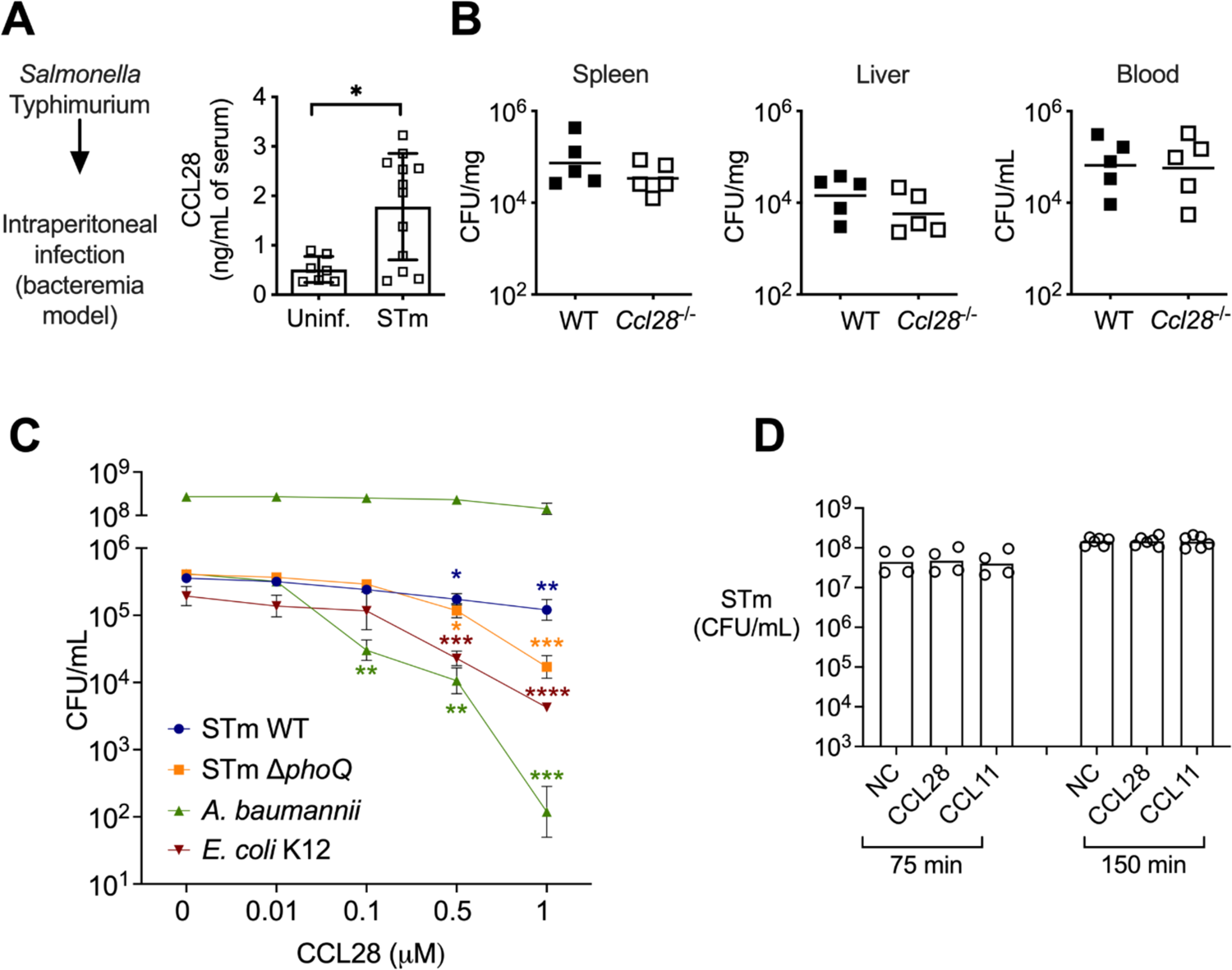
CCL28 does not confer protection in a *Salmonella* bacteremia model, and lacks direct antimicrobial activity against *Salmonella*. (A, B) For the bacteremia model, mice were infected by intraperitoneal injection with *S.* Typhimurium (STm, 1×10^3^ CFU) or sterile PBS (uninfected control). (A) At 96h post-infection, CCL28 in serum was quantified by ELISA of wild-type mice (uninfected, n=7; STm, n=12). Data shown comprise two independent experiments. Bars represent the mean ± SD. (B) STm CFU was determined in the spleen, liver, and blood of WT mice (black squares) and *Ccl28^-/-^* mice (white squares) 96h after intraperitoneal infection with STm (1×10^3^ CFU). Data shown comprise two independent experiments (WT, n=5; *Ccl28*^-/-^, n=5). (C, D) *In vitro* antimicrobial activity of CCL28 against STm wild-type, STm *ΔphoQ*, *E. coli* K12, and *A. baumannii*. (C) 5×10^5^ CFU/mL of each strain (*A. baumannii* additionally at 5×10^8^ CFU/mL) was incubated with recombinant murine CCL28 at the indicated concentrations (n=4 per group), and CFU were enumerated after 2h. (D) STm wild-type (1×10^7^ CFU/mL) was incubated with recombinant murine CCL28 (50 nM) or CCL11 (25 nM) and CFU were enumerated at 75 min (n=4 per group) and 150 min (n=6 per group). Bars represent the geometric mean. (A) Data were analyzed by Mann-Whitney U relative to uninfected controls. (C) Log-transformed data were analyzed by nonparametric one-way ANOVA (Kruskal-Wallis) for independent samples. Dunn’s multiple comparison test was performed to compare bacterial CFU at each time point relative to time zero (control group). Significant changes are indicated by \**p* ≤ 0.05, \*\**p* ≤ 0.01, \*\*\**p* ≤ 0.001 \*\*\*\**p* ≤ 0.0001.

**Supplementary Figure 2.**
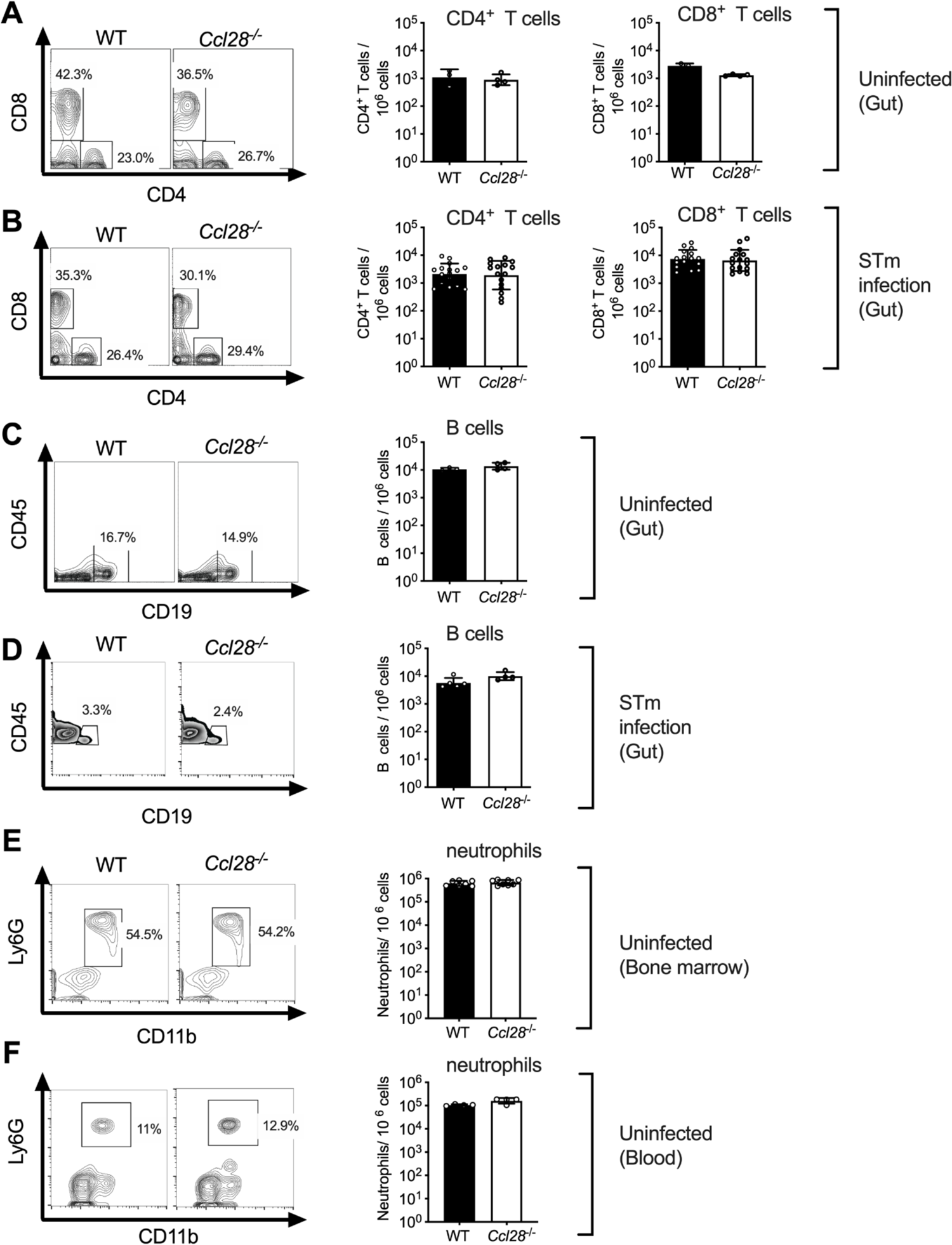
Wild-type and *Ccl28^-/-^* mice exhibit similar numbers of T and B cells, as well as bone marrow and blood neutrophils. Flow cytometry analysis of (**A**, **B**) CD4^+^ and CD8^+^ T cells, and (**C, D**) CD19^+^ B cells isolated from the gut of (**A, C**) uninfected WT and *Ccl28^-/-^* mice or (**B, D**) WT and *Ccl28^-/-^* mice infected with STm for 48h (colitis model; see also Fig. 1). (**E, F**) Cells from (**E**) bone marrow or (**F**) blood of uninfected WT (black bars) or *Ccl28^-/-^* (white bars) mice were analyzed by flow cytometry to determine the percentage and number of neutrophils. (**A-F**) Left panels show representative contour plots. Right panels show the frequency of the indicated cells per million CD45+ live cells of WT mice (black bars) and *Ccl28^-/-^* mice (white bars). Each circle represents a mouse. Bars represent the geomean ± SD.

**Supplementary Figure 3.**
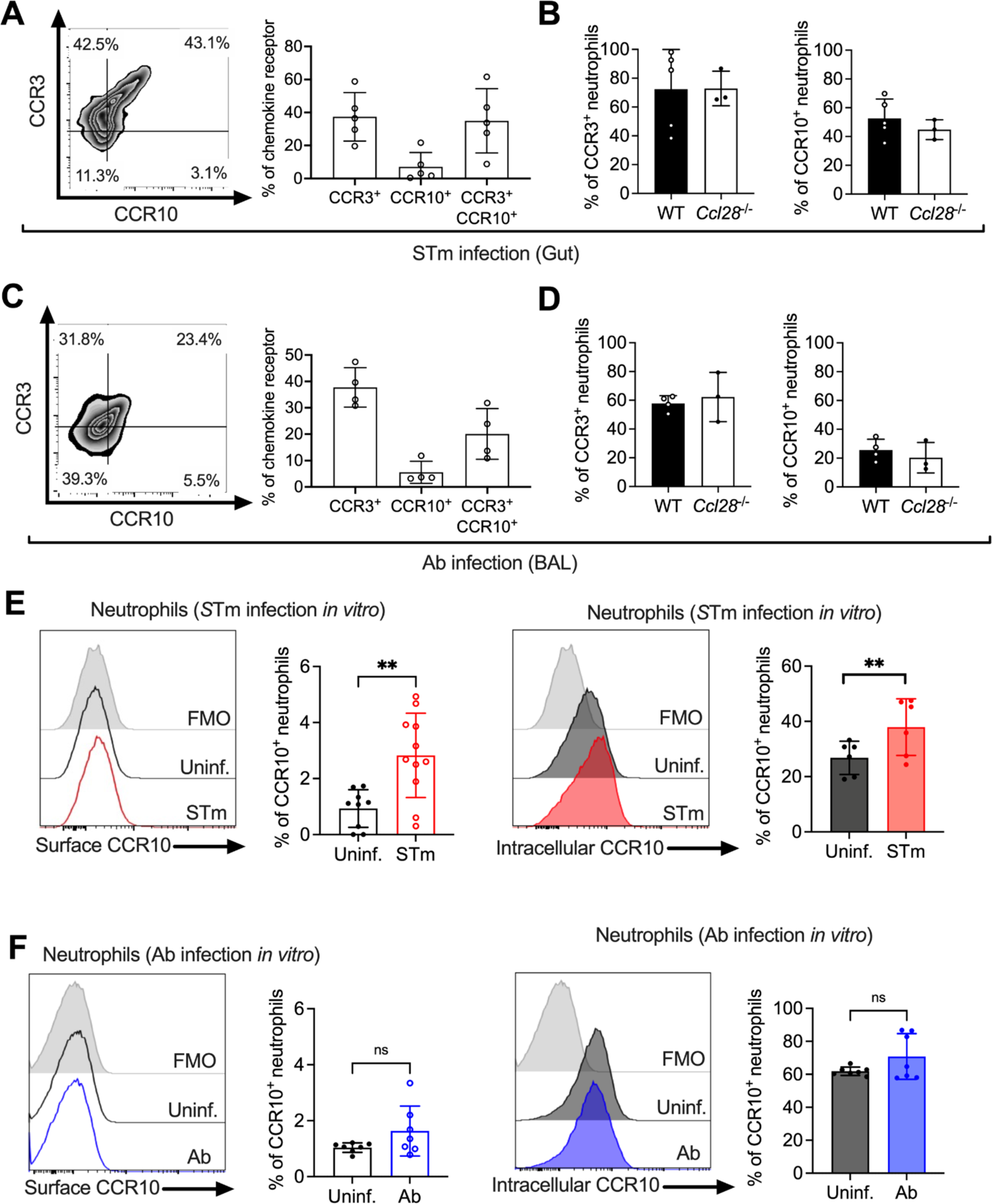
Expression of CCR3 and CCR10 in neutrophils isolated from the gut and lung mucosa, and CCR10 in neutrophils infected *in vitro*. (**A**) Surface expression of CCR3 and CCR10 on neutrophils obtained from the gut of WT mice (n=5) infected with STm for 72h, analyzed by flow cytometry. (**B**) Percentage of CCR3^+^ and CCR10^+^ neutrophils obtained from the gut of WT (n=5) and *Ccl28*^-/-^ mice (n=3) infected with STm for 72h, analyzed by flow cytometry. (**C**) Surface expression of CCR3 and CCR10 on neutrophils obtained from the BAL of WT mice (n=4) infected with Ab for 24h, analyzed by flow cytometry. (**D**) Percentage of CCR3^+^ and CCR10^+^ neutrophils obtained from the BAL of WT (n=4) and *Ccl28*^-/-^ mice (n=3) infected with Ab for 24h, analyzed by flow cytometry. (**A, C**) Left panels show representative contour plots, and right panels show the percentages of neutrophils expressing the indicated receptor on their surface. Bars represent the mean ± SD. (**E, F**) Bone marrow neutrophils were infected with (**E**) opsonized STm at MOI=10 for 1h or (**F**) Ab at MOI=10 for 4h. Surface (clear histograms) or intracellular (solid histograms) CCR10-expressing neutrophils were detected by flow cytometry. Left panels show representative histograms, and right panels show the percentage of neutrophils expressing CCR10 on their surface (clear bars) or intracellularly (solid bars). (**E, F**) A significant change (paired *t* test) relative to uninfected controls is indicated by \*\**p* ≤ 0.01; ns = not significant.

